# A gate and clamp regulate sequential DNA strand cleavage by CRISPR-Cas12a

**DOI:** 10.1101/2021.06.18.448962

**Authors:** Mohsin M. Naqvi, Laura Lee, Oscar E. Torres Montaguth, Mark D. Szczelkun

## Abstract

CRISPR-Cas12a has been widely used for genome editing and diagnostic applications, yet it is not fully understood how RNA-guided DNA recognition activates the sequential cleavage of the non-target strand (NTS) followed by the target strand (TS). Here we used single-molecule magnetic tweezers microscopy, ensemble gel-based assays and nanopore sequencing to explore the coupling of DNA unwinding and cleavage. In addition to dynamic R-loop formation, we also directly observed transient dsDNA unwinding downstream of the 20 bp DNA:RNA hybrid and, following NTS cleavage and prior to TS cleavage, formation of a hyperstable “clamped” Cas12a-DNA intermediate resistant to DNA twisting. Alanine substitution of a conserved aromatic amino acid “gate” in the REC2 domain that normally caps the heteroduplex produced more frequent and extended downstream DNA breathing, a longer-lived twist-resistant state, and a 16-fold faster rate of TS cleavage. We suggest that both breathing and clamping events, regulated by the gate and by NTS cleavage, deliver the unwound TS to the RuvC nuclease and result from previously described REC2 and NUC domain motions.

---

The clustered regularly interspaced short palindromic repeats (CRISPR)/CRISPR-associated (Cas) nuclease effectors such as Type II-A Cas9 and Type V-A Cas12a that evolved as prokaryotic defence systems against bacteriophage have found widespread utility in gene editing (1–3). There is evidence that Cas12a has superior properties as an engineering tool by having fewer off-target binding and cleavage events (4–6). Additionally, Cas12a has been exploited for nucleic acid recognition diagnostics, including detection of SARS-CoV-2 genomic RNA (7). In both systems, RNA-guided DNA recognition occurs by strand separation of a protospacer target to allow Watson-Crick base pairing between the DNA targeted-strand (TS) and the spacer sequence of a CRISPR RNA (crRNA), and the release of a non-targeted strand (NTS) (8). Type II enzymes such as Cas9 employ two nuclease domains with distinct phylogenies, RuvC and HNH, to cut the NTS and TS, respectively (9,10). In contrast, Type V CRISPR effectors have a single RuvC domain which for the prototype Cas12a (formerly known as Cpf1) cuts DNA in an obligatory sequential mechanism; NTS cleavage first followed by ∼20-fold slower TS cleavage (11–16). However, it is unclear how the Cas12a RuvC domain transitions between cleaving the two strands (17). A better understanding of this mechanism will aid in the design of Cas12a enzymes with improved properties for gene editing and/or diagnostic applications.

Cas12a structures are bi-lobed monomer, consisting of recognition and nuclease lobes (Figure 1a) (12,13,18–23). The Cas12a-crRNA complex first scans a DNA target until the flexible pocket formed by the wedge (WED), REC1 domain and PAM-interacting domain (PID) interacts with a specific protospacer adjacent motif (PAM, 5′-TTTV-3′, where V = A/C/G). Upon DNA binding, the PID reduces its amplitude of motion, while REC2 and Nuc motions are increased so that they move outward to help accommodate the TS in the RuvC active site (13,22–24). PAM distortion leads to ATP-independent stand separation and R-loop formation within the recognition lobe, starting the pre-structured 3′ end of the crRNA spacer sequence (the “seed”) (15,20,22,25). As R-loop formation progresses, conformational checkpoints need to be passed to activate DNA cleavage (13,20,26,27): a loop connecting REC1 and REC2 lobes (the “linker”) firstly interacts with the 5th to 7th nucleotides of the crRNA; a loop (the “lid”) then interacts with the 8th to 11th crRNA nucleotides; and finally, a REC1 helix (the “finger”) interacts with the 15th to 17th crRNA nucleotides; additionally, the bridge helix plays a role in stringency and coordination of conformation changes. At least 17 bp of hybrid is required to satisfy the checkpoints and activate cleavage and Cas12a binding and cleavage is tolerant of mismatches in the final 3 bp of the R-loop (6,12,16,25,28,29). Nonetheless, high resolution crystal structures show complete 20 bp R-loops forming with pseudo A-form structure (e.g. Figure 1a, (21)).

**Figure 1:**
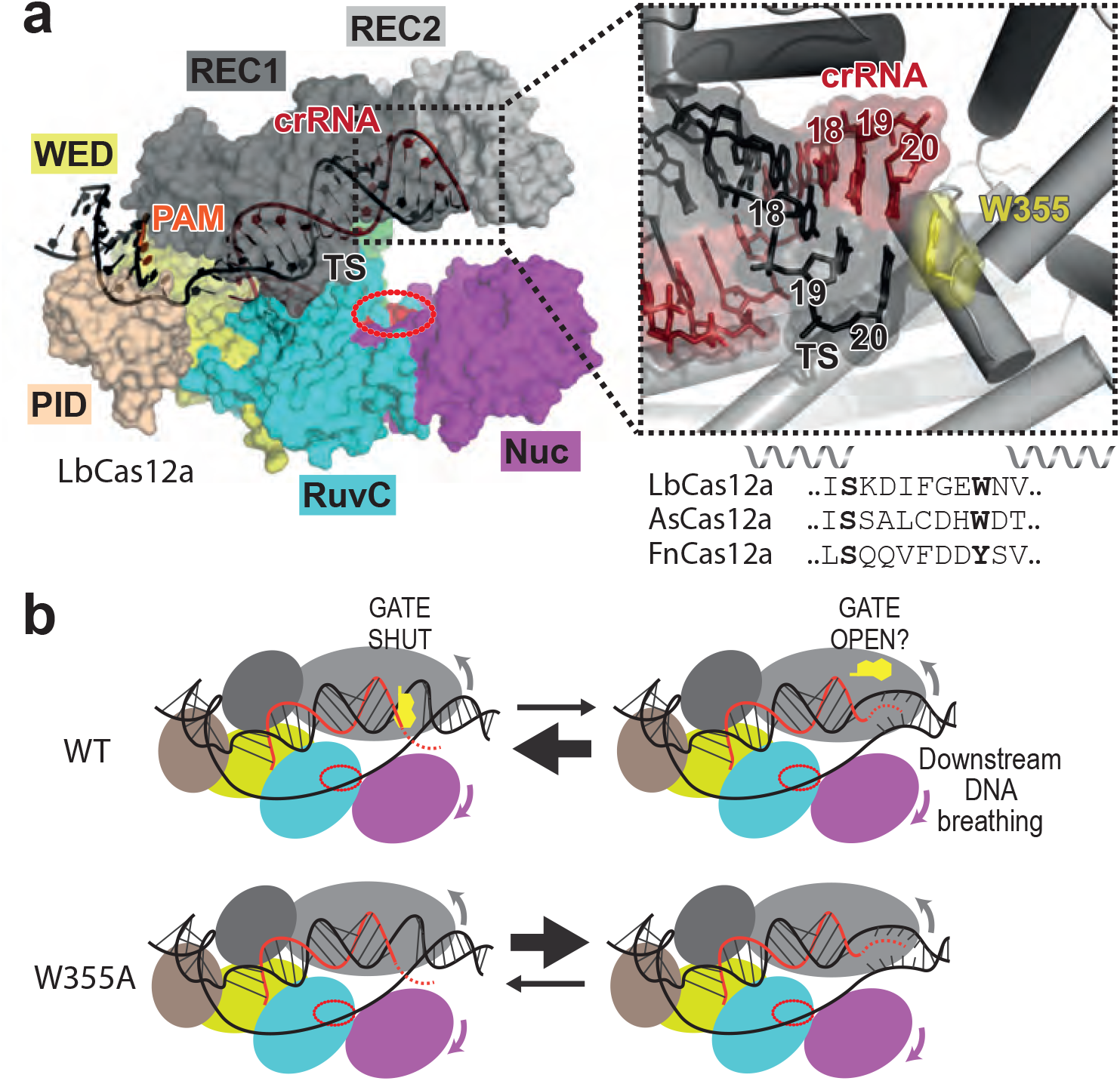
The Cas12a aromatic stacking Gate and downstream DNA breathing. (**a**) Structure of LbCas12a (PDB:5xus, (21)) with REC1 (dark grey), REC2 (light grey), Wedge (WED, yellow), Pam Interaction Domain (PID, wheat), RuvC nuclease (cyan) with catalytic site (red oval) and NUC (magenta) shown as protein surface, and DNA (black) and crRNA (red) as cartoons. The path of the NTS is not resolved in this structure. The inset shows the structure around the aromatic gate (W355 in LbCas12a) that stacks against the 20^th^ base pair of the R-loop between the crRNA and TS. The amino acid sequence alignment of the Gate loop is shown for LbCas12a, *Acidaminococcus sp*. BV3L6 Cas12a (AsCas12a) and *Francisella novicida* Cas12a (FnCas12a). (**b**) Cartoon representation of the REC2/Nuc domain motions and opening of the Gate that supports PAM-distal DNA breathing. A W355A mutant would have the Gate permanently open, favouring DNA breathing states.

Following R-loop formation the liberated NTS docks into the RuvC active site and nicking occurs (Supplementary Figure S1). The DNA ends then disengage and an uncut NTS phosphodiester can re-enter the active site and rounds of cleavage-release-rebinding lead to DNA end trimming (30). This gap formation may prevent steric occlusion of the open active site to allow subsequent TS binding and allow binding of non-specific ssDNA *in trans* (“bystander” cleavage), a key activity for nucleic acid detection diagnostics (31). A geometric limitation for TS cleavage is that the target site is displaced (32), in a region of dsDNA downstream of the R-loop, ∼25 A° from the RuvC active site and in an incorrect orientation for in-line nucleophilic attack without rotation to match the polarity of the NTS (8,11). A suggested model is that downstream dsDNA unwinds, and the released single stranded TS is bent towards RuvC (Supplementary Figure S1) (13). Structures consistent with these molecular gymnastics have been observed for the related Cas12b and Cas12f complexes (33,34). Cryo-EM structures, single molecule FRET measurements and molecular dynamics of Cas12a additionally support an inward closing motion of the REC2 and Nuc domains following NTS nicking (13,24). Slower TS cleavage may reflect both the kinetics of TS gap formation and these domain transitions.

More recently, Cofsky et al (30) demonstrated that the downstream DNA that encompasses the TS cleavage site is subject to DNA breathing. They propose this is a fundamental property of the RNA 3′ end of an R-loop that is exploited by Type V enzymes due to their R-loop geometry. Extending the region of downstream ssDNA accelerated DNA cleavage and altered the TS cleavage loci, confirming the important role of the downstream unwinding and the kinetic constraint it places on the TS cleavage rate. Here we were able to corroborate these findings in the absence of DNA cleavage by directly observing the dynamics of transient and reversible downstream unwinding events by *Lachnospiraceae bacterium ND2006* (Lb) Cas12a using a magnetic tweezers assay that can measure strand separation in real time (15,35). We also observed that the LbCas12a R-loop was highly dynamic and heterogeneous, forming multiple intermediate stable states and occasionally fully unzipping (rupture events).

It has been suggested that REC2 dynamics could control conformational changes in Nuc, regulating a functional role in loading the TS into RuvC (12,33). Cas12a ternary X-ray structures reveal that the R-loop is terminated by a stacking interaction with a conserved aromatic amino acid in REC2 (e.g., W355 in LbCas12a, Figure 1a, (21)). Although mutation of this residue had moderate reduced INDEL formation activity (20), we speculated that this “Gate” may control the downstream DNA breathing by transitioning between closed (stacked) and open states (Figure 1b). Indeed, mutation of the W355 Gate to alanine resulted in more frequent and extended downstream DNA breathing but also greater occupancy of longer R-loop states in general and less state heterogeneity, ruling out a role for the stacking interaction in stabilising the 20 bp R-loop. Using ensemble endonuclease assays, we demonstrated that the more frequent DNA breathing with W355A accelerated the rate of endonucleolytic TS cleavage by ∼16-fold. Subsequent 5′-3′ processing of the cleaved end was also accelerated as mapped by nanopore sequencing. By following nuclease activity in the single molecule tweezers assay, we also observed a torque-resistant clamping of the downstream DNA after NTS cleavage, which we suggest corresponds to Nuc interactions that guide the TS to the RuvC active site. This clamped state was hyperstable in the W355 mutant and supported more rapid TS cleavage. We therefore propose that the aromatic Gate residue acts to control both downstream DNA breathing and the clamping that is necessary for TS cleavage.

## RESULTS

### Single molecule observation of dynamic R-loops and downstream DNA breathing

To observe Cas12a-dependent DNA unwinding events in real time, we used a single molecule magnetic tweezers assay used previously with cas9 and Cas12a (Figure 2a) (15,35). A 2 kbp linear DNA target was tethered between a glass coverslip and a magnetic particle (500 nm diameter) in a flow cell. The apparent length of the DNA above the surface was monitored by video microscopy of the bead image (36). A pair of permanent magnets above the flow cell stretched the DNA with varying force depending on the vertical position and could be rotated to introduce positive or negative turns into the DNA, inducing supercoiling at low forces. We initially used an EDTA-based buffer so that the dynamics of R-loop formation could be monitored without DNA cleavage (15).

**Figure 2:**
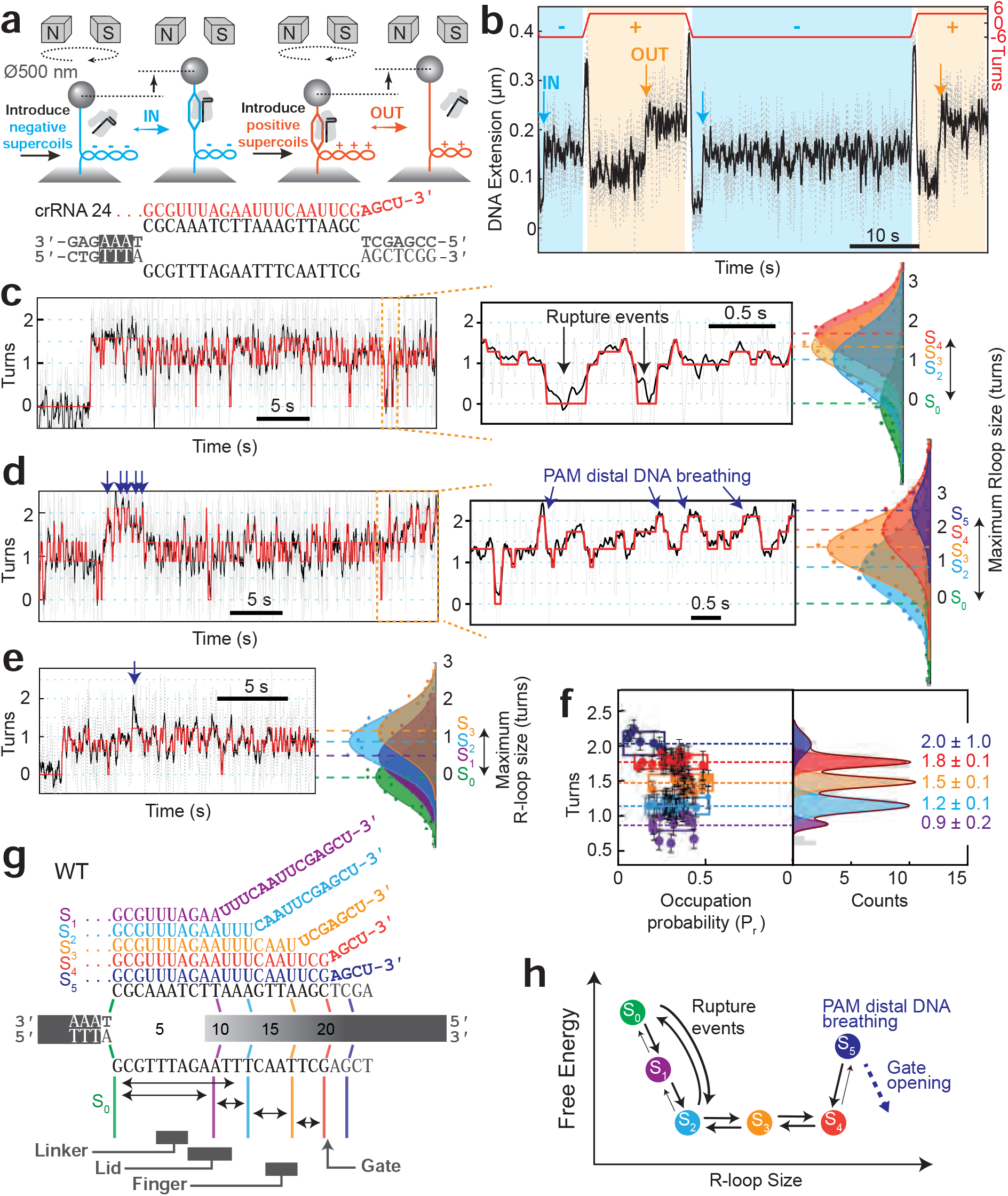
Wild Type Cas12a produces dynamic R-loop states. (**a**) Representation of the magnetic tweezers experiments and DNA protospacer and crRNA spacer sequences used (see main text). (**b**) An example extension time trace (for all data, grey 60 Hz raw data, black 10 Hz filtered) at 0.3 pN showing R-loop formation (IN, blue, at -7 pN nm) and dissociation (OUT, orange, at +7 pN nm) events by WT LbCas12a. (**c**) Example R-loop formation trace at -7 pN nm measured as changes in turns showing hopping between 4 states identified by HMM fitting (red); R-loop states S_2_, S_3_, S_4_ and reversible transitions to an R-loop dissociated state (rupture events), S_0_. Gaussian fitting of the histograms gives the average position of each state in agreement with the HMM fitting. (**d**) Example R-loop formation trace at -7 pN nm showing hopping between 5 states including an extended state (blue arrows, S_5_) that represents pam-distal breathing. (**e**) Example R-loop formation trace at -7 pN nm showing hopping between 4 states, including an additional short R-loop (S_1_). (**f**) Plot showing the 5 R-loop states identified by HMM analysis and their probability of occupation (P_r_) from multiple traces (*N*=34), with state positions confirmed from Gaussian fitting. Box width corresponds to the full width at half maximum of each peak. P_r_ Error bars are SD in the turn positions measured by HMM. Errors bars in turn values are s.d from the Gaussian peak fitting. (**g**) Cartoon representation of the R-loops for each state and relative positions of structural features. (**h**) Simple free energy diagram showing sequential hopping between R-loop states. Opening of the Gate residue would lower the energetic barrier to the S_5_ state.

At the 0.3 pN stretching force used here, introducing negative turns formed negative supercoils that shortened the apparent DNA length (Figure 2b). Negative torque supports R-loop formation (15), which resulted in a reduction in DNA supercoiling to balance the altered DNA linking difference, observed as an increase in apparent DNA extension (IN events in Figures 2a,b). To force the R-loop out, positive turns were introduced to generate positive supercoiling. The positive torque causes R-loop dissociation (OUT events in Figures 2a,b). The process was repeated by cycling between negative and positive turns using the magnets. Hereafter we used a linear relationship to convert DNA extension to DNA turns (Supplementary Figure S2) so that we could compare between different DNA molecules where relative attachment points can cause variations in apparent DNA length.

An example R-loop formation profile at -7 pN nm using 1 nM WT Lb Cas12a is shown in Figure 2c. The improved signal-to-noise using the smaller diameter (500 nm) magnetic particles (37), revealed a much more dynamic R-loop than was detectable previously. Each trace was fitted with a Hidden Markov model (HMM) to identify the minimal number of discrete states that could describe the data and to calculate the state positions in turns and their relative probabilities. The example data in Figure 2c could be described by 3 discrete states (S_2_, S_3_, and S_4_) showing sequential hopping transitions. We noted that there were frequent “rupture” events where the turns returned to zero, suggesting a complete reversible unzipping of the R-loop (S_0_).

A second example profile (Figure 2d) illustrates the heterogeneity in R-loop dynamics, since the S_3_ state is more occupied and we could identify an additional state, S_5_, with a turns value of 2.0 ± 1.0. We interpret the transient S_5_ events as PAM distal DNA breathing, as observed by Cofsky et al (see below) (30). A third example profile (Figure 2e) reveals another discrete state (S_1_) representing a much shorter R-loop. The full length S_4_ R-loop state does not appear to be accessed during this event, although there is a transient increase that could be accessing S_4_ and/or S_5_, but this could not be identified from the HMM analysis because of its infrequency during the measurement window. In general, there was great heterogeneity between events, both in the total turn sizes, number of states, and dynamics of the states (Supplementary Figure S3).

The occupation probability (P_r_) and average turn size of the states (Figure 2f) and the rupture event probability (Figure 3e) were calculated from multiple events (Supplementary Figure S3). We previously measured that the 20 bp R-loop of Cas12a corresponds to a change of 1.8 turns (15). This matches the average turns for state S_4_ here. We therefore mapped the other states onto possible R-loop sizes (Figure 2g). From the occupation probabilities, we can also suggest a simple free energy diagram for the different states (Figure 2h). The S_1_, S_2_ and S_3_ states could correspond to the previously identified conformation checkpoints that couple R-loop propagation to nuclease activation [8]. The linker and lid interactions may produce the occupancy of the S_1_ and S_2_ states, respectively. HMM modelling suggests that the rupture events can occur directly from these states. The “finger” interaction may produce the occupancy of the S_3_ state.

**Figure 3:**
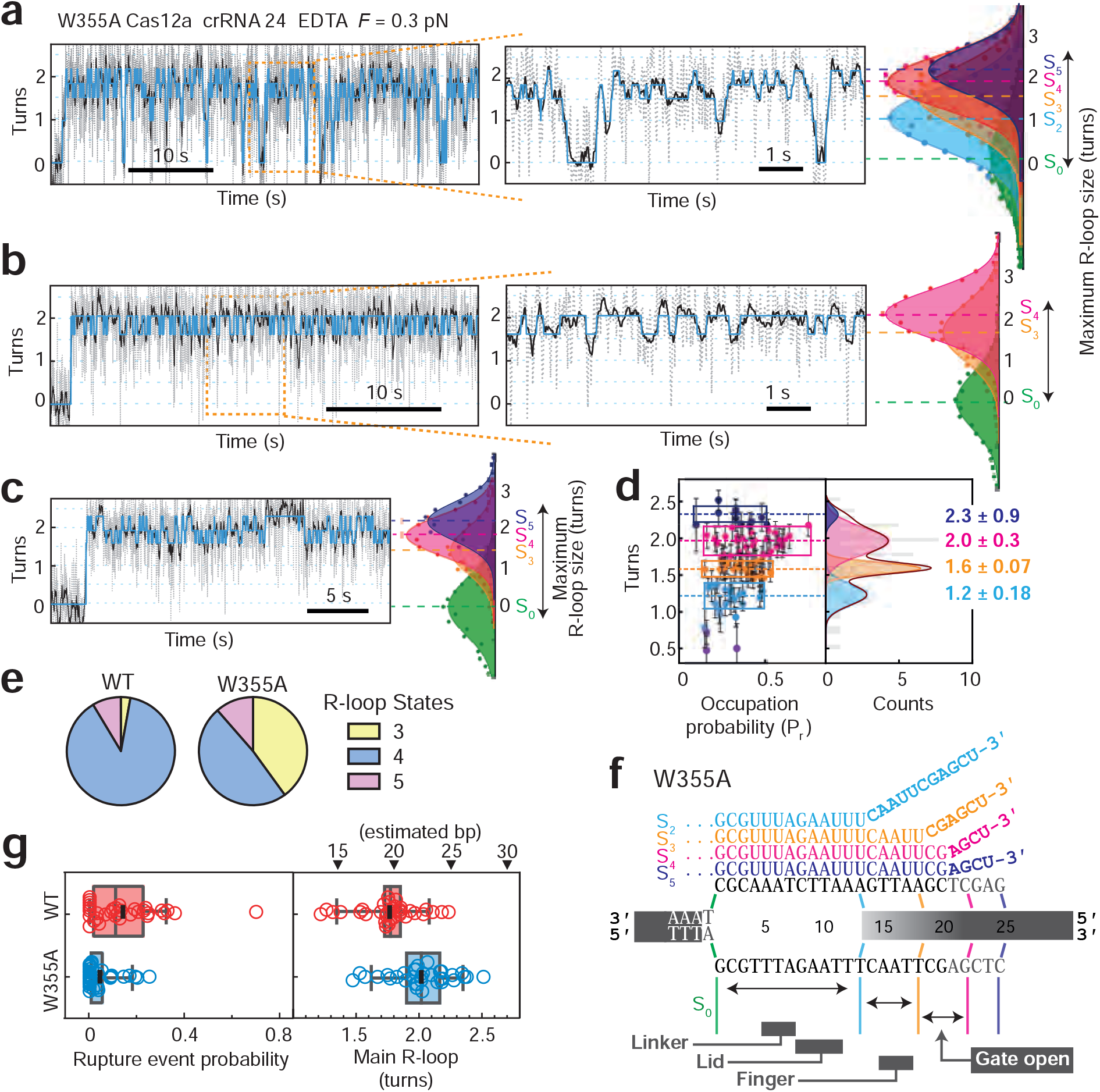
Mutation of the Gate residue stabilises downstream DNA breathing and longer R-loop states, and reduces the frequency of R-loop rupture. (**a**) Example R-loop formation trace at -7 pN nm with W355A Cas12a showing hopping between 5 states identified by HMM fitting (blue); S_2_, S_3_, S_4_ and S_5_, and reversible rupture events to S_0_. Gaussian fitting of the histograms gives the average position of each state in agreement with the HMM fitting. (**b**) Example R-loop formation trace at -7 pN nm showing hopping between only 2 states; S_3_ and S4. The slower R-loop formation at the start allowed clear identification of S_0_ without rupture occurring. (**c**) Example R-loop formation trace at -7 pN nm showing hopping between 3 states; S_3_, S4 and S_5_. As in (b), rupture events were not observed but the initial S_0_ could be identified. (**d**) Plot showing the 4 R-loop states identified by HMM analysis and their probability of occupation (P_r_) from multiple traces (*N*=34), with state positions confirmed from Gaussian fitting. Box width corresponds to the full width at half maximum of each peak. P_r_ Error bars are SD in the turn positions measured by HMM. Errors bars in turn values are the s.d from the Gaussian peak fitting. State S_4_ (pink) shows maximum occupation. The downstream DNA breathing events (in blue) show more stability and frequency than observed for Wt Cas12a (Figure 2f). (**e**) Pie charts of percentage of traces showing 3, 4 or 5 states for Wt and W355A Cas12a events. (**f**) Cartoon representation of the R-loops for each state. (**g**) Box plots comparing rupture event probability and main R-loop size measured from HMM analysis for WT (red, N=34) and W355A (blue, N=34) Cas12a.

The S_5_ state (2.0±1.0 turns) can be estimated to correspond to an additional unwinding of ∼2 bp. Although we cannot rule out that the change in turns partly corresponds to a change in writhe, this additional unwinding is consistent with the observations of Cofsky et al (30). We suggest there is a free energy penalty to accessing this state from the full R-loop S_4_ state due to the W355 Gate (see Discussion). The transient downstream unwinding may be due to movement (“opening”) of the Gate residue that otherwise caps the R-loop. This idea is tested in the next section.

In summary, our data reveals a more dynamic R-loop than was presumed previously, rapidly switching between S_2_-S_4_ states and even a fully ruptured R-loop state, possibly controlled by reversibility of the conformational checkpoints. An additional S_5_ state is consistent with a transient and reversible unwinding of a few additional base pairs downstream of the R-loop.

### The aromatic Gate controls downstream DNA breathing and favours longer R-loop states

To test whether the W355 residue of LbCas12a stabilises the R-loop and/or regulates the frequency of downstream DNA breathing events, we mutated the residue to an alanine to mimic a putative open conformation (Figure 1b) and tested the R-loop dynamics as above. Example events are shown in Figures 3a-c. In general, W355A demonstrated a greater occupancy of longer R-loop states (Figure 3d, Supplementary Figure S3) and there were fewer states identified by the HMM fitting per event (Figure 3e). The S_2_ state was less frequently observed than with WT, while the S_1_ state was not occupied to a measurable degree (Figure 3d). The most occupied state was 2.0±0.3 turns (S_4_) which we suggest corresponds to a 20 bp R-loop plus 2 further unwound base pairs, equivalent to the transient S_5_ state of the WT enzyme (Figure 3f). In other words, the Gate mutant seems to lock in the additional downstream breathing. The additional occupied S_5_ state corresponds to 2.3±0.9 turns, which we interpret as additional downstream unwinding by 2-3 bp not observed as a long-lived state with WT. Rupture events were observed, but less frequently that with WT (Figure 3g), which may reflect that the shorter R-loop states were less occupied and so there was less chance for R-loop collapse.

We previously used rapid switching to positive supercoiling to measure the R-loop lifetime under varying positive torque. Our observations of dynamic states here suggest that it would be difficult to reliably capture a defined S-state and that there may also be dynamic changes during the magnet rotation period (∼1 s). Furthermore, although stable states at positive torque were noted for WT (Figure 2b), as observed previously (15), the majority of W355A events were immediately dissociated so that lifetime distributions could not be measured (Supplementary Figure S4). This result was surprising given that rupture events at negative torque were infrequent and the average R-loop states were longer in length. However, if R-loop rewinding under positive torque is driven from the PAM-distal end (the PAM end being clamped by the PID), the difference in stability may reflect that in the WT enzyme, the Gate can resist the rewinding force.

In summary, the removal of the Gate residue resulted in more stable R-loops and additional downstream unwinding at negative torque, but unstable R-loop formation at positive torque. The change in turns captured was greater than that with WT, indicating that the downstream DNA was breathing more readily (Figure 3h). These results suggest that stacking of W355 against the terminal bases of the heteroduplex does not stabilise formation of a 20 bp R-loop but does control the unwinding/rewinding at the PAM distal end. An open state of the Gate therefore favours downstream breathing but is also more sensitive to rewinding force.

### The aromatic Gate limits the rate of target strand cleavage and subsequent nucleolytic processing of the cleaved ends

If downstream breathing is necessary to start the processes of docking the TS into the RuvC active site (30), we reasoned that the stabilisation of downstream unwound states observed with W355A (Figure 3) would accelerate the TS cleavage rate. To measure DNA cleavage, we collected time points from an ensemble time course using a supercoiled plasmid DNA substrate (SC) where the nicked intermediate (Open Circle, OC) and cleaved linear product (LIN) can be separated by agarose gel electrophoresis and the bands quantified (Figure 4a) (15). The cleavage profiles for WT and W355A Cas12a were fitted by numerical integration (Figure 4b), to return the rate constants for ordered cleavage of the NTS followed by TS (Figure 4c). As observed previously (15), the WT enzyme cuts the TS >20-fold slower than the NTS (Figure 4b,c). In contrast, W355A cleaved the TS at almost the same rate as the NTS (Figure 4a-c). This result is consistent with an activation of the second cleavage event resulting from the increased downstream breathing when the Gate is absent (Figure 3). The NTS cleavage rate were only slightly slower for W355A compared to WT, so the altered R-loop dynamics and enhanced DNA breathing of the gate mutant did not have a major effect on the first strand cleavage.

**Figure 4:**
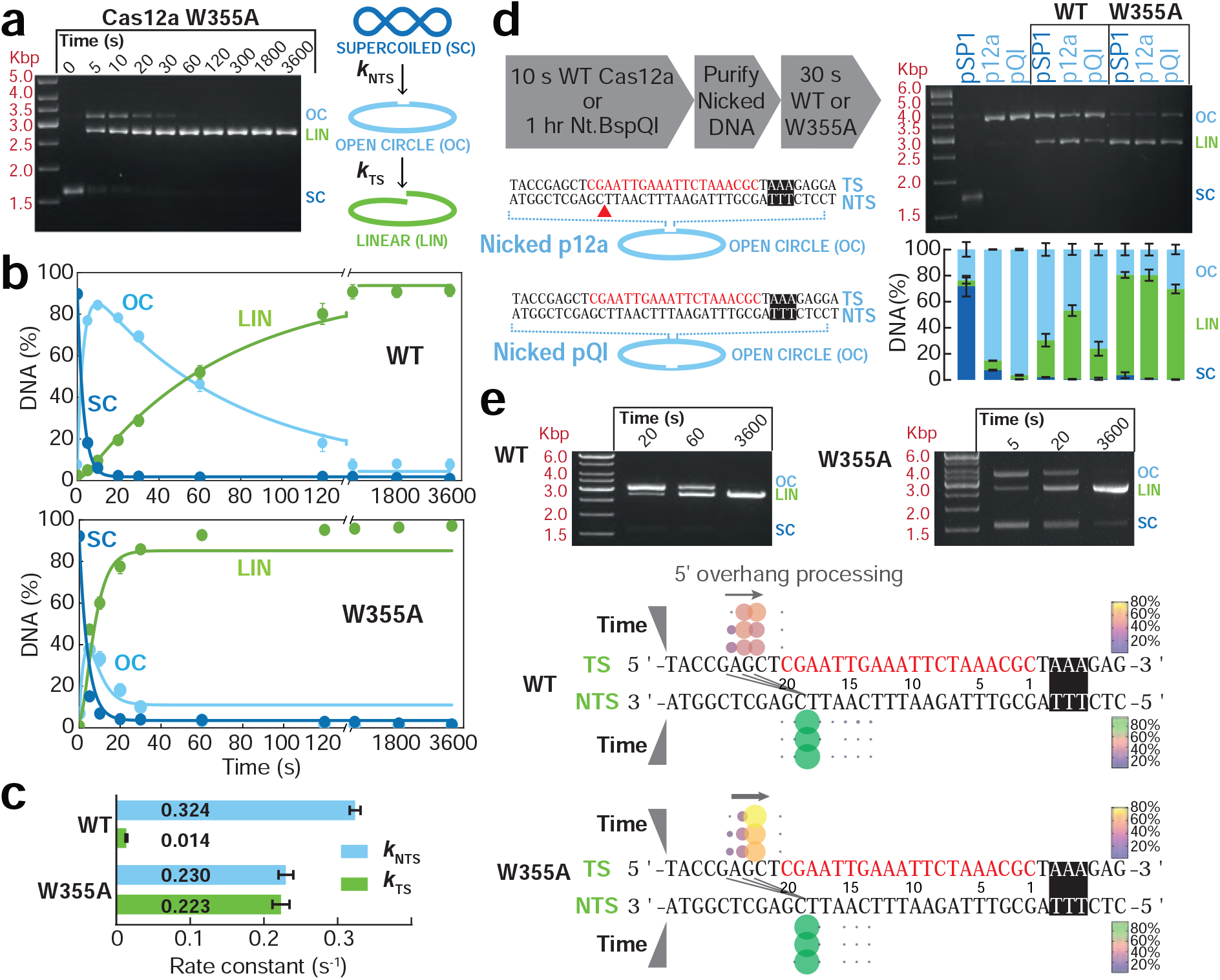
Mutation of the Gate residue accelerates endonucleolytic cleavage of the targeted strand. (**a**) Agarose gel of pSP1 cleavage time course using W355A Cas12a. Diagrams represent the supercoiled substrate (SC), nicked (open circle) intermediate (OC) and linear product (LIN) states, with the rate constants *k*_NTS_ and *k*_TS_ representing the ordered cleavage of the NTS followed by TS. (**b**) Quantified data from individual cleavage time courses with WT or W355A Cas12a were simultaneously fitted using numerical integration to a simple cleavage model and the rate constants averaged (15). Solid circles are means from 3 repeats while solid lines represent model simulations using the average parameters (in c). (**c**) Mean rate constants for cleavage of the NTS and TS from 3 repeats (error bars, SD). (**d**) The effect of nicked DNA on relative NTS and TS cleavage. Plasmid DNA was pre-treated with either Cas12a (to nick the NTS within the protospacer, red triangle) or Nt.BspQI (to nick at a distal site). The purified nicked DNA was treated with WT or W355A Cas12a for 30 s and the DNA separated by gel electrophoresis. Average percentages of SC, OC and LIN from 3 repeats were quantified by gel densitometry (errors bars, SD). (**e**) Cleavage reactions using pSP1 and either WT or W355A Cas12a were stopped at the times shown and the cleaved ends of linear DNA from individual dsDNA cleavage events mapped using ENDO-Pore (38). Circle diameter and colour both represents the relative percentage cleavage at that location on each strand (*N*: 3064, WT 20 s; 5812 WT 60 s; 5597 WT 3600 s; 3063 W355A 5 s; 2604 W355A 20 s; 3984 W355A 5 s). Three diagonal lines represent the linkage between NTS and TS cleavage events for >92% of events. W355A produces the same cleavage loci as WT but the 5′-3′ processing of the TS strand is faster (grey arrows). Note that ENDO-Pore returns cleavage loci of a single event that are closest to the 3′ end of each strand regardless of the order of cleavage (Supplementary Figure S7).

The steps leading to engagement of the TS by RuvC may be triggered by the breakage of the NTS strand or may have a requirement for RuvC to go through its cleavage chemistry in a more ordered conformational exchange. To test these alternatives, we took advantage of the slow TS cleavage rate of Cas12a to produce a pre-nicked DNA substrate by 10 s treatment with WT enzyme (Figure 4d). Since the cleavage rate of nicked DNA is rate-limited by the slower R-loop formation compared to supercoiled DNA, a control pre-nicked DNA was produced using Nt.BspQI cleavage at a distal site (Figure 4d). The cleavage of the nicked substrates after 30 s using either WT or W355A Cas12a was compared with cleavage of supercoiled DNA (Figure 4d). For WT enzyme, pre-nicking of the NTS resulted in more cleavage of the TS, indicating that a barrier had been lifted. In contrast, W355A produced similar levels of linear DNA on all substrates, consistent with the removal of the Gate having a larger effect on accelerating the cleavage process.

Given that the locations of NTS and TS cleavage can be altered by mismatches within the R-loop (29) and that RuvC-dependent processing of the DNA ends can also be affected by downstream DNA unwinding (30), we reasoned that the differences in R-loop dynamics and DNA breathing between WT and W355A Cas12a might also result in measurable differences in cleavage loci. To explore this, we used ENDO-Pore, a nanopore-based method that allows the linked-end mapping of single endonuclease DNA cleavage events with median site accuracy of >99.5% (38). DNA cleavage libraries were generated from the linear DNA generated by WT or W355A reactions at three time points (Figure 4e), and then sequenced and analysed (Materials and Methods, Supplementary Figure S6). WT Cas12a produced the expected staggered 5′ cleavage. More than 95% of NTS cleavage events were located at position 18, although there is evidence of low levels of 5′ cleavage events that are consistent with the main product resulting from 5′-3′ processing of NTS cleavage events that occurred prior to the earliest time points collected. On the TS, at each time point there were three principal cleavage loci, at positions 22, 23 and 24, producing 5′ single strand overhangs of either 4, 5 or 6 nt, respectively (Figure 4e). There was a gradual shift from the 6 nt to 4 nt overhang with time, providing further evidence of slow 5′-3′ TS processing following initial cleavage (30).

Using W355A, the positions of the main cleavage events were the same as WT except that the 5′-3′ processing of TS was noticeably faster, with the majority product being the shortest 4 nt overhang at the earliest time point (5 s).

RNA-guided TS hybridization conformationally activates the Cas12a RuvC domain, triggering a non-specific, single-stranded DNase activity that can target DNA *in trans* (31). The resulting “bystander” DNA cleavage has been exploited for Cas12a diagnostic tools as the activity is catalytic and thus provides an amplification of any RNA-guided recognition of a nucleic acid. Although W355A has an activated nuclease activity on the TS, it had a slower and less efficient bystander cleavage activity compared to WT Cas12a (Supplementary Figure S5). This suggests that although the removal of the Gate allows quicker access to conformations necessary for TS cleavage, the nuclease activity of W355A is not overactive *per se*, or that the mutant protein adopts a conformational state where the RuvC active site is less accessible to *in trans* DNA.

In summary, removal of the W355 aromatic gate results in faster endonucleolytic and exonucleolytic DNA cleavage of the TS. We suggest that the activated downstream DNA unwinding may be more quickly delivering the TS to the nuclease domain. Cofsky et al also noted that destabilisation of the DNA downstream of the R-loop resulted in faster TS cleavage Since the positions of DNA cleavage remain the same, the overall structural geometry of the complex is likely to be unaltered.

### A stable downstream DNA Clamp state following non-target strand cleavage

To further explore the nuclease mechanism and role of the W355 Gate, we adapted the magnetic tweezers assay by using a Mg^2+^-based buffer that supports DNA cleavage but kept all other conditions the same. A DNA was positively supercoiled before enzyme was introduced into the flow cell; under positive torque, R-loop formation is inhibited so the DNA is not cut. The DNA was then rapidly unwound (<1 s) to produce negatively supercoiled DNA (event 1 in Figure 5a), facilitating R-loop formation recorded as a change in apparent bead height (event 2). We expected that R-loop formation would then support NTS cleavage and that free rotation at the NTS nick would release the negative supercoiling strain, producing a further increase in apparent bead height to full length (39). Such an event was observed (event 3). Because of the limited time resolution of the assay, we could not identify whether this event was preceded by a DNA breathing event (i.e., state S_5_). Subsequent TS cleavage would produce a DSB, breaking the bead-DNA tether which would be observed as loss of bead tracking. However, we noted unexpected properties of the nicked intermediate that are explained below and in a model in Figure 5b.

**Figure 5:**
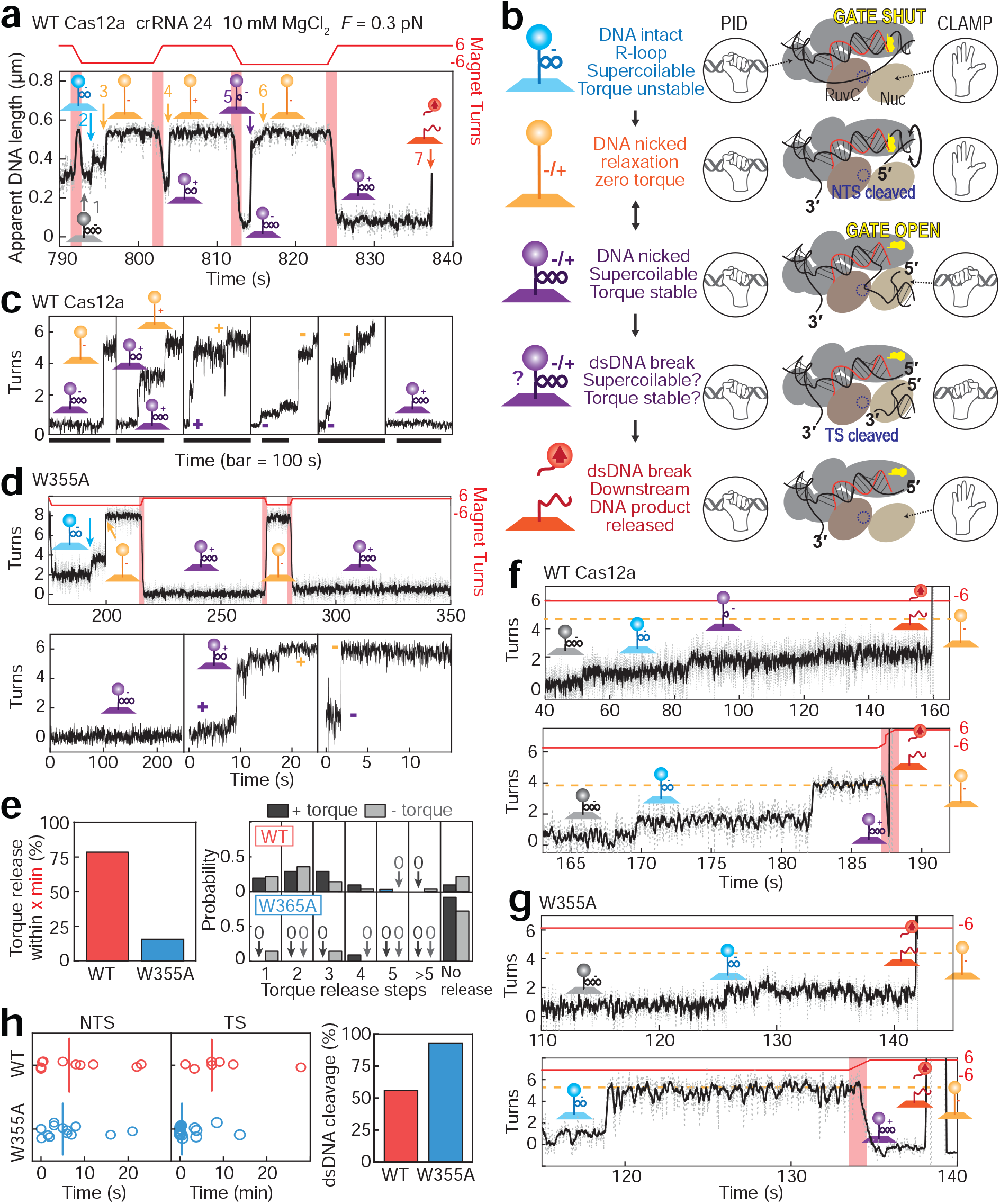
Cleavage of the non-targeted strand by Cas12a results in clamping of downstream DNA controlled by the Gate to produce a torque stable state. (**a**) Example extension time trace (grey 60 Hz raw data, black 10 Hz filtered) at 0.3 pN showing R-loop formation and DNA cleavage with WT Cas12a and Mg^2+^ ions. Magnet rotations from positive to negative values are shown in red. Numbered events are explained in the main text. The cartoons are also explained in panel b. (**b**) Steps in the cleavage pathway. Cas12a is shown in cartoon form as in Figure 1b. The effective topology of the DNA-bead tethers is shown in the cartoons, left. The hand cartoons represent clamped and unclamped states of the PID and, nominally, the NUC. The W355 Gate (yellow) is shown shut (vertical, stacked against R-loop) or open (horizontal). See main text for full explanation. (**c**) Example WT traces showing nicked DNA in torque stable clamped states (purple cartoon) and nicked, relaxed states (orange) at positive (+7 pN nm) or negative (−7 pN nm) torque. Note the heterogeneity in kinetics and number of turns released per step during unclamping. Rightmost panel shows an example of stable clamping over 100 s. (**d**) (*upper panel*) Example W355A trace showing R-loop formation (blue), torque release following NTS nicking (orange) and subsequent positive torque-stable clamped states (purple). (*lower panel*) Example W355A traces comparing nicked DNA in typical torque stable clamped states (purple) and rare torque release steps producing relaxed states (orange). (**e**) (*left panel*) Percentage of clamped events at positive or negative torque that released (N=13/63 for WT, N=15/18 for W355A, *p* < 0.05). Note for W355A, most clamped events resulted in TS cleavage (see panel g). (*right panel*) Number of release steps to produce a relaxed state from the clamped state at +7 pN nm (blue) or -7 pN nm (grey). Where the clamped state did not release, the average waiting time was 93 ± 20 s. (**f**) Example WT traces showing dsDNA cleavage. (*upper graph*) Following R-loop formation (blue), the DNA is partially relaxed into a clamped state (purple) until bead tracking is lost due to TS cleavage event (red). (*lower graph*) Following R-loop formation (blue), the DNA is relaxed at NTS cleavage (orange). Loss of bead tracking due to TS cleavage (red) is observed from a clamped state during magnet rotation. (**g**) Example W355A traces showing dsDNA cleavage. (*upper graph*) Following R-loop formation (blue), bead tracking is lost due to dsDNA cleavage (red) without obvious release to a relaxed or clamped state. (*lower graph*) Following R-loop formation (blue), the DNA is relaxed at NTS cleavage (orange). Loss of bead tracking due to TS cleavage (red) is observed from a clamped state during magnet rotation. (**h**) Scatter plots for WT (red) and W355A (blue) showing time between R-loop formation and DNA relaxation (NTS cleavage, *left panel*) and between DNA relaxation and loss of bead tracking (TS cleavage, *middle panel*). Lines are medians of distributions. (*right panel*) Bar plot showing percentage of molecules showing NTS and TS cleavage for WT (N = 8/14) and W355A (N= 16/17) (*p* < 0.05). For DNA that only showed NTS cleavage, TS cleavage was not observed after an average of 26 minutes.

Following the increase in bead height interpreted as resulting from NTS cleavage (event 3 in Figure 5a), the tethered DNA should have been nicked and thus mechanical rotation of the magnets should not have had an effect. However, introducing twelve positive turns resulted in a lowering of the bead height to an apparent DNA length that represents ∼7 trapped positive supercoils after ∼0.5 s. Following a delay of ∼0.5 s, the apparent DNA height increased to full length in a single step (event 4), consistent with rapid free rotation at the nick (40). After a delay, twelve negative turns were introduced by magnet rotation, and again a reduction in bead height was observed. The shortening happened immediately and produced a final apparent DNA length consistent with 11-12 trapped negative supercoils. Following a delay of ∼1 s, the apparent DNA height increased to close to full length in a rapid step (event 5). The remaining trapped negative supercoils were more gradually released over ∼2 s to produce full length relaxed DNA (event 6). Finally, introduction of twelve positive turns produced a positively supercoiled DNA (12 trapped supercoils). After a delay at ∼11.5 s, a loss of bead tracking (event 7) indicated a DSB in the DNA-bead tether.

These observations can be explained by a model where Cas12a transiently clamps the downstream DNA following NTS cleavage to trap the R-loop and DNA strand breakages in an isolated topological domain (Figure 5b). In the initial R-loop formed at negative torque, the PAM proximal DNA end is clamped by the PID and cannot rotate whereas the PAM distal downstream DNA is free to rotate. Hence the change in bead height in event 2 (Figure 5a) as the change in twist resulting from R-loop formation can dissipate throughout the DNA. We propose that the Gate is “shut” when a full 20 bp R-loop forms. Upon NTS cleavage by RuvC, the downstream DNA can freely rotate, relaxing the DNA and reducing the torque to zero. NTS cleavage also triggers opening of the Gate, which allows reversible unwinding of the downstream DNA. This conformation change in REC2 is coupled to inward motion towards Nuc and DNA clamping. We suggest that these interactions are with the Nuc domain. The broken NTS is then trapped within an isolated topological domain by the PID and Nuc clamps. Consequently, twisting of the DNA by magnet rotation can trap supercoils in the rest of the DNA. This double-clamped state is in dynamic equilibrium with the single-clamped state, possibly by opening-closing transitions of the Gate, so trapped supercoiling may again release by free rotation at the nick after a delay. In the double-clamped state, the TS is held close to the RuvC active site. If cleavage occurs, then the double-clamping prevents immediate product release. Once the Nuc clamp releases the DNA, the downstream product will be released. This is consistent with previous observations that Cas12a retains interaction with the PAM-proximal product and releases the PAM distal DNA (25).

An important consequence of our model is that because both DNA relaxation following NTS cleavage and bead loss due to TS cleavage will only be observed if the Nuc clamp is open, observed lifetimes of the R-loop state and the full-length state, respectively, do not necessarily correspond solely to the chemical strand breakage rates but could also reflect the stochastics of the clamp opening and closing. Nonetheless, we can test the model by comparison between WT and W355A Cas12a. In W355A, the Gate is permanently open, so we predicted that the Nuc clamp state would be more readily accessed and thus the NTS nicked state would be more torque stable. This stable clamped state would also explain the faster TS cleavage (Figure 4a-c), since this state helps the TS engage with the nuclease active site (Figure 5b).

Shown in Figure 5c are a series of examples of NTS nicked states that have been put under negative (−7 pN nm) or positive (+7 pN nm) torque by magnet rotation and have trapped supercoils. Supercoiling release occurred in steps, the size and lifetime of which varied stochastically. In 79.4% of cases, full supercoiling release was observed within a 93 s observation window (Figure 5e), while in cases where no release was observed, 75% showed TS cleavage. The median number of release steps was 2 regardless of the sign of the torque (Figure 5e). In comparison, W355A was more torque stable following NTS cleavage (Figure 5e). In 83.4% of cases, this state was not released and in 88% of such cases led directly to the loss of bead tracking, indicating TS cleavage and clamp release (Figure 5b). In the other 16.6% of cases, release events were observed which showed stepped release similar to WT but with insufficient number of events to reliably determine the median number of steps. We interpret this data to show that for WT, the Gate can reversibly shut, conformationally triggering the clamp to release the trapped supercoils. The clamp is less likely to release in W355A where the Gate is always in an open state.

The slower TS cleavage rate for WT observed in the ensemble cleavage assays (Figure 4c) may be due to the clamped state being less occupied. Complete examples of R-loop formation and dsDNA cleavage are shown for WT and W355A in Figures 5f and 5g, respectively. In both upper panels, events are shown where following NTS cleavage, there is only partial release of supercoiling, consistent with the clamp closing during relaxation at the nick. TS cleavage and loss of bead tracking occurs from this clamped state. In both lower panels, events are shown where following NTS cleavage, there was full relaxation of the supercoils. In these examples bead loss due to TS cleavage was observed following magnet rotation and trapping of supercoils. Similar to these examples, in most cases for both WT and W355A, cleavage occurred from identifiable clamped states, as mentioned previously.

The times for NTS cleavage estimated from the single molecule assays were similar for WT and W355A (Figure 5h). This suggests that the observed difference in clamping occurs after TS cleavage; i.e., initial R-loop lifetimes reflect the chemical cleavage kinetics. As observed in the ensemble reactions, TS cleavage measured from bead release was faster for W355A, although there were far fewer measurable events for WT Cas12a over a reasonable experimental timescale (see below). Bead loss with WT Cas12a was only observed in 57% of molecules over the experimental timescale while for W355A this value was significantly higher at 94 % (*p < 0*.*05*) (Figure 5h). The less frequent cleavage by WT could be due to the less stable clamped state (Figure 5e), or to TS cleavage being inhibited by applied force, possibly by preventing conformation changes needed for gate opening.

In summary, single molecule cleavage experiments revealed that Cas12a can form a clamped state following NTS cleavage that is substantially more torque stable than the R-loop formed in EDTA. The stability of the clamped state is promoted by opening of the Gate and appears to be necessary for TS cleavage. Consequently, the slow TS cleavage of WT Cas12a can be accelerated by removing the Gate in the W355A mutant.

## DISCUSSION

We explored the R-loop dynamics and activation of NTS and TS cleavage by Cas12a using a combination of single molecule R-loop assays, ensemble DNA cleavage assays, single cleavage event mapping by nanopore sequencing (ENDO-Pore) and single molecule cleavage assays. Our results show that an aromatic residue in the REC2 domain that terminally caps the full 20 bp heteroduplex, which we term the Gate (Figure 1a), does not stabilise the R-loop directly through stacking interactions but controls downstream DNA breathing (Figure 1b), and following NTS cleavage, conformational activation of a clamped state that is necessary for primary TS cleavage and subsequent end processing (Figure 5b).

By using smaller diameter magnetic particles in our magnetic tweezers assays, we could observe dynamic interchange between defined R-loops states in the absence of DNA cleavage that could be mapped to approximate locations of previously identified conformational checkpoints (Figures 2g & 3f). We also captured reversible R-loop rupture events where the DNA briefly fully rewound and DNA breathing states downstream of the R-loop that are likely to be important for TS cleavage (Figures 2 & 3) (30). The pathway of R-loop formation was heterogenous between different events (Supplementary Figure S3). Removal of the Gate (W355A) produced fewer transitions between states and favoured longer R-loops, suggesting that the stacking of the aromatic amino acid against the terminal bases does not stabilise the 20 bp R-loop, and its absence actually produces fewer transitions between states. A kinetic analysis of AsCas12a was also consistent with readily reversible R-loop propagation until formation of a full 20 bp R-loop and with R-loop formation limiting the cleavage rate (14).

A key difference between Type II-A and Type V-A CRISPR-Cas effectors is that the former use two separate nuclease domains to cut the NTS and TS while the latter use a single RuvC domain that must transition sequentially between the NTS and TS (8). It has been suggested that R-loop asymmetry is exploited by Cas12a to allow downstream DNA breathing that is a key step in providing the single strand TS that can be delivered to the RuvC nuclease active site (30). We directly corroborated such reversible DNA breathing events for WT Cas12a as transient increases in DNA unwinding measured from changes in supercoiling (Figure 2 & 3). Additionally, we found that in the W355A mutant there was an increase in frequency and size of the breathing events (Figure 3), resulting in an increased rate of TS cleavage and processing (Figures 4 & 5).

Our results are compatible with the idea of asymmetry in the R-loop (30) since the role of the Gate in stacking against the end of the R-loop may be to buffer the effect. i.e. the stacking of an aromatic amino acid prevents breathing until strand separation is needed. Breathing may only occur if the Gate opens transiently, opens due to the trigger of NTS cleavage, or is artificially removed by mutation. The rate of TS cleavage is therefore partly restricted by the Gate opening (and thus DNA breathing) and partly by the necessity for NTS cleavage and gap formation (Figure 4d) (30).

When measuring DNA cleavage in our single molecule assay (Figure 5), we observed post-NTS cleavage states that could stably trap DNA torque. For WT Cas12a these were transient states that released the trapped supercoils within seconds. However, for the Gate mutant the states were torque-stable over tens of seconds, an unexpected result given that we rarely observed positive torque stable states in the absence of NTS cleavage. The increased clamped state lifetime corresponded with a higher frequency of TS cleavage during the experimental observation window, consistent with the ensemble assays in Figure 4. The release of the clamped state appeared instantaneous, suggesting that there is not a frictional control of DNA rotation as observed for Type IB topoisomerases (40); instead the clamp exists in binary open and closed states.

We interpret the clamped state as downstream protein-DNA interactions that are necessary for delivering the TS to the RuvC active site (13,24,30). Hence TS cleavage is produced in a clamped state. Our data is consistent with clamping being controlled by the Gate since its removal favoured the clamped state and hence accelerated the observed TS cleavage rate. The reversibility of the clamping seen with WT Cas12a could be due to reversible opening-closing of the Gate even following NTS cleavage. The 5′-3′ processing of the TS (Figure 4e) is also favoured by the Gate mutant. Clamping may therefore also play a role in repeatedly delivering the cleaved 5′ end of the TS into the RuvC active site (so that both NTS and TS are subject to gap formation). In contrast, the bystander activity was reduced with W355A (Supplementary Figure S5; possibly the efficient formation of the clamp state inhibits interaction of the RuvC active site with DNA *in trans*.

Single molecule, structural and molecular dynamics studies of Cas12a have indicated that the REC2 and Nuc domains move towards each other upon NTS cleavage, contracting the groove between the TS and RuvC active site (12,13,24). We interpret the clamped state observed after NTS cleavage (Figure 5), as resulting from this motion. Since a combined path created by residues from both RuvC and Nuc domains is essential for TS cleavage (12,33), we further suggest that TS interactions by Nuc, possibly following DNA breathing, are responsible for the torque stable clamped state (Figure 5b). Equivalent auxiliary target nucleic acid-binding (TNB) domains that help load the TS into the RuvC active site are found across the Type V family [e.g. the Nuc domain for Cas12a and Cas12b or, the target-strand loading (TSL) domain for Cas12e, (34,41)]. Although they adopt distinct structures (42), a general role for TNBs could be to act as a Clamp necessary for TS delivery to the RuvC active site (12,34). Based on their proposed roles and coordination, engineering of TNB and/or Rec2 domains may be a fertile ground for producing new Type V enzymes with improved DNA cleavage properties, either by reducing off-target cleavage by rejecting TS cleavage or by favouring more open states that are necessary for bystander cleavage.

## METHODS

### Protein production and ribonucleoprotein assembly

The W355A LbCas12a mutation was generated by overlap extension PCR (primers 5′-AGTAAAGACATTTTCGGTGAGGCGAACGTGATCCGTGACAAATGG-3′ and 5′-CCATTTGTCACGGA TCACGTTCGCCTCACCGAAAATGTCTTTACT-3′) with pSUMOCas12a (15). WT and W355A LbCas12a were expressed and purified as published previously (15). Based on additional breakdown products, W355A appears less stable than WT enzyme (Supplementary Figure S8). crRNAs were synthesised and HPLC-purified by IDT (5′-UAAUUUCUACUAAGUGUAGAUGCGUUUAGAAUUUCAAUUCGAGCU-3′). For Ribonucleoprotein (RNP) complex assembly, 250nM Cas12a and 250nM crRNA were mixed in buffer RB [10 mM Tris-Cl, pH 7.5, 100 mM NaCl, 10 mM MgCl_2_, 0.1 mM DTT, 5 μg/ml BSA] supplemented with 1 U per 20 µL SUPERase-In RNase Inhibitor (ThermoFisher) and incubated at 37 °C for 1 hour.

### Single Molecule Magnetic Tweezers Experiments

The magnetic tweezers experiments were performed using a commercial PicoTwist microscope (Fleurieux sur L’Arbresle, France) equipped with a 60 Hz Jai CV-A10 GE camera (36). For flow cell preparation, glass coverslips (Menzel Gläser No.1, 24 × 60 mm x160 µm) were cleaned in 3 repeated cycles of 1 hr sonication in 1M KOH and then Acetone and were subsequently cleaned with milliQ water and dried using compressed air. The coverslips were kept enclosed in glass jars to prevent any moisture until needed. Flow channels were prepared as previously. DNA molecules (a 2 kbp section of pSP1) were tethered to 500 nm paramagnetic beads (Adamtech)(37), and the glass coverslip of the flow cell as previously described (15,35). Topologically-constrained DNA were identified from rotation curves at 0.3 pN and the rotational zero reference (Rot_0_) set. 1 nM RNP was used for all measurements. The R-loop formation and dissociation experiments (Figures 2 & 3) were performed in Buffer SB [10 mM Tris-Cl, pH 7.5, 100 mM NaCl, 1 mM EDTA, 0.1 mM DTT, 5 μg/ml BSA] at 25 °C while the single molecule cleavage experiments (Figure 5) were performed in Buffer RB at 25 °C.

### Hidden Markov modelling of R-loop sizes and dynamics

Individual R-loop formation traces were sorted out from the raw data (60 Hz) using custom built Matlab code and were filtered to 10 Hz. The DNA extensions were converted into turn values by linear fitting of the constant torque region of each hat curve (10 turns/s) at negative torque (Supplementary Figure S2). Each trace was then fitted with a Hidden Markov model (HMM) (43). From long time series of observations, HMM determines the kinetics between different states defined by a transition matrix and the gaussian signal of each state, that assign each point in the observation sequence the most likely state of the system at a given time (44). Both with WT (Figure 2) and mutant Cas12a (Figure 3) traces, the best fitting HMM was determined by extracting the turn values for each state in the HMM model using a custom-built Fortran code. The histograms of turn values of each state were then separately fitted with a gaussian model (in Origin Lab) to extract the state positions. The HMM that described state positions separated by > 0.1 turns was considered as the best fit as lower values resulted in the merging of Gaussian peaks and poor fitting of the traces. Next, for each trace the occupation probability (P_r_) of the states was determined by calculating the ratio of the lifetimes of each state (using HMM) and the total time of the measured R-loop event. The rupture event probability was measured by counting the number of events showing transitions from the R-loop to basal state and dividing by the total time of the trace. The MATLAB and Fortran codes used for analysis are available upon request.

### Statistical Analysis

The statistical significance of differences in torque release percentages (Figure 5e) and dsDNA cleavage percentages (Figure 5h) between experimental conditions was calculated using one tailed two proportion z-test. Test results are mentioned as *p* values in the legends. In box charts, whiskers indicate 90% and 10% extreme values, the inner line represents the median, the length of the box indicate interquartile range and the black small vertical bar the mean of the population.

### Ensemble DNA cleavage assays

5 nM tritiated pSP1 was pre-heated in Buffer RB at 25 ºC for 5 minutes. Reactions were started by addition of 50 nM Cas12a RNP and incubated for the time specified. The reaction was quenched by adding 0.5 volumes of STEB [0.1 M Tris (pH 7.5), 0.2 M EDTA, 40% (w/v) sucrose, 0.4 mg/ml bromophenol blue] and incubating at 67 ºC for 10 minutes. Samples were separated by agarose gel electrophoresis on a 1.5 % (w/v) agarose gel in 1X TAE [40 mM Tris-acetate, 1 mM EDTA, 10 μg/ml ethidium bromide] at 2 V/cm overnight (16 hours) and visualised by UV irradiation. DNA bands containing supercoiled, linear or open circle DNA were excised and placed into scintillation vials. 0.5 ml sodium perchlorate was added to each gel slice, and tubes were incubated at 67 ºC for 2 hours to melt the agarose. The vials were cooled to room temperature and 10 ml Hionic-Fluor Scintillation Cocktail (Perkin Elmer) added to each vial and shaken thoroughly. Each vial was counted in a Tri-Carb Trio 3100TR Liquid Scintillation Counter for 10 minutes. Where indicated, the cleavage data was fitted to the model described in Mullally et al using numerical integration in Berkeley Madonna (www.berkeleymadonna.com).

To make pre-nicked DNA, 5 nM tritiated pSP1 was nicked either using 50 nM Cas12a RNP (WT protein with crRNA 24) at 25°C in Buffer RB for 10 s or using 10 units Nt.BspQI (New England Biolabs) per µg DNA, at 50°C for 1 hr. Nicked DNA was purified using Qiagen PCR purification kit. Cleavage assays were then performed as described above and stopped after 30 s by addition of 0.5 volumes of STEB followed by incubation at 67 ºC for 10 minutes.

For the bystander assays (Supplementary Figure S5), 50 nM RNP (WT or W355A with crRNA 24) was preincubated with 5 nM ssDNA activator (5′-GGATCCCCGGGTACCGAGCTCGAATTGAAATTCTAAACGCTAAAGAGGAAGAC-3′) for 30 min at 37°C. Negative RNP controls were incubated for 10 min in the absence of ssDNA activator. M13mp18 ssDNA (New England Biolabs) was added to 5 nM, and reaction aliquots quenched using STEB at the time points indicated. Samples were separated by agarose gel electrophoresis on a 1.5 % (w/v) agarose gel in 1X TAE at 2 V/cm overnight (16 hours), and post-stained with ethidium bromide (0.5 ug/ml). The intact M13 band was quantified by densitometry (ImageJ) from a low exposure image. High exposure images were collected to clearly show the smear more clearly in Supplementary Figure S5.

### ENDO-Pore linked end mapping of Cas12a DNA cleavage

Cleavage reactions using RNP (WT and W355A with crRNA 24) and pSP1 were quenched at different time points, as above, and the DNA purified using a DNA Clean & Concentrator-25 kit (Zymo Research). End-repair and dA tailing were performed using NEBNext® Ultra− II End Repair/dA-Tailing Module (New England Biolabs) and the DNA ligated with a dT tailed chloramphenicol cassette, recording the cleavage position. OmniMAX− 2 T1R *E. coli* cells (ThermoFisher) were transformed with the ligation reactions and single colonies selected using 34 μg/mL chloramphenicol. Cleavage event libraries were generated by scrapping and pooling >20,000 colonies followed by plasmid purification. Rolling circle amplification (RCA) was then performed using 10 ng of the cleavage library as template with EquiPhi29− DNA Polymerase and exonuclease resistant random hexamers (ThermoFisher). Reactions were incubated for 2 hours at 45°C, heat inactivated for 10 min at 65°C, and the DNA purified using AMPure XP beads (Beckman Coulter). RCA products were debranched using 10 units/ µg DNA of T7 Endonuclease I (New England Biolabs) for 15 min at 37°C. The debranching reaction was stopped by incubating with 0.8 units of Proteinase K (New England Biolabs) for 5 min at 37°C. Debranched products were purified using AMPure XP beads followed by size selection using the Short Read Elimination XS kit (Circulomics). Samples were prepared for Nanopore sequencing using the Ligation Sequencing Kit (SQK-LSK109) combined with the Native Barcoding Expansion kit and sequenced using R9.4.1 cells (Oxford Nanopore Technologies). Circular concatemeric sequences were generated using C3POa v2.2.2 (45). Individual dsDNA breaks on pSP1 were identified using sequences with ≥5 concatemer repeats using bespoke software (Cleavage Site Identifier) (Supplementary Figure S6) (38).

## Supporting information

Supplementary results

## ACKNOWLEDGEMENTS

We thank Stephen Cross for help with ENDO-Pore software development. This work was support by the BBSRC (BB/S001239/1) and the European Research Council under the European Union’s Horizon 2020 research and innovation programme (ERC-2017-ADG-788405).

## AUTHOR CONTRIBUTIONS

M.D.S. conceptualized the study. L.L. purified recombinant proteins. M.M.N. and M.D.S. designed all single-molecule assays. M.M.N. performed and analyzed data from single-molecule assays. L.L. and M.D.S. designed the ensemble cleavage assays. L.L. performed and analyzed data from ensemble cleavage assays. O.E.T.M. designed the ENDO-Pore sequencing. L.L. and O.E.T.M. performed and analyzed data from the ENDO-Pore sequencing. All authors contributed to the original draft and reviewed and edited the manuscript.

