## Supplementary results for "A gate and clamp regulate sequential DNA strand cleavage by CRISPR-Cas12a"

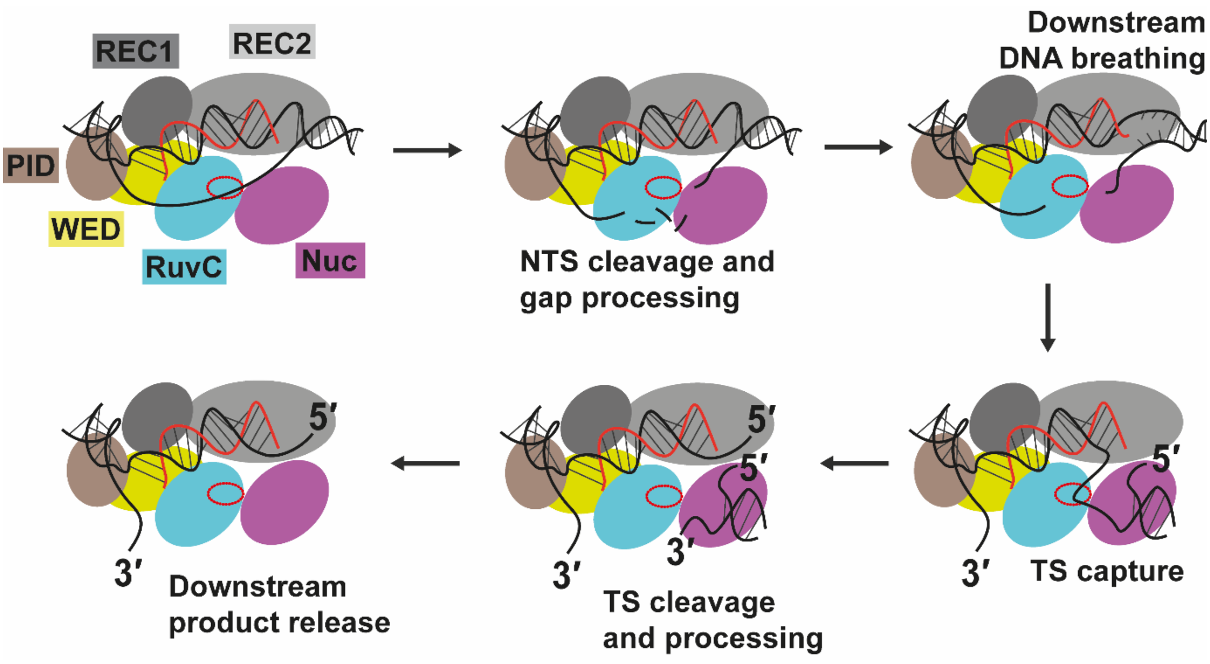


**Supplementary Figure S1. Model for sequential cleavage of the NTS and the TS by Cas12a.** Cartoon representation of Cas12a subdomains and conformation changes necessary for sequential dsDNA cleavage. The RuvC active site is shown as a red oval. See main text for full details.

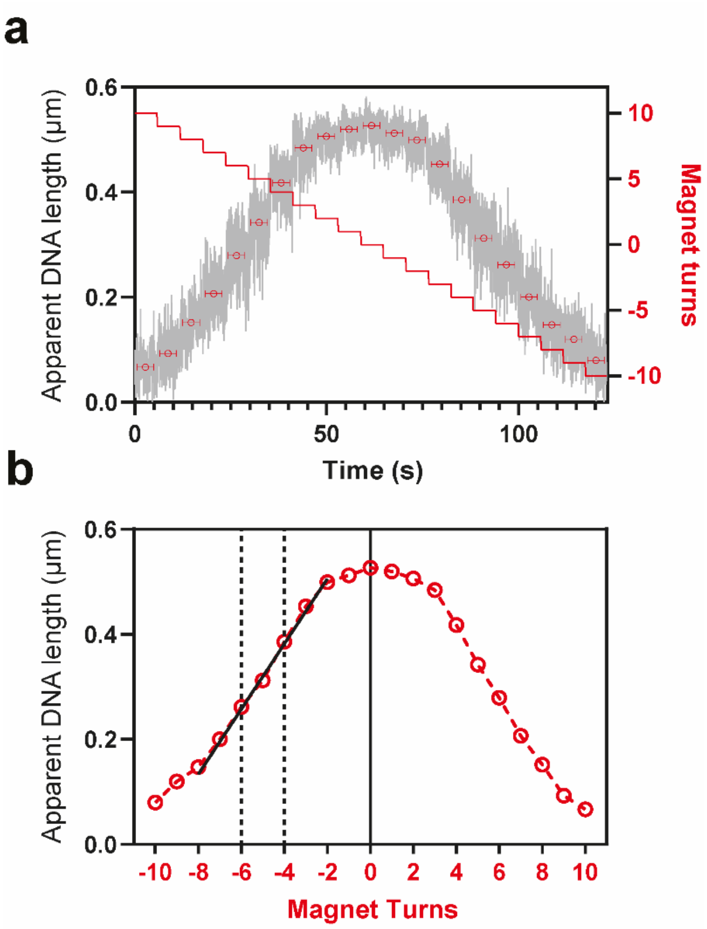


**Supplementary Figure S2. DNA rotation-extension curve (Hat curve).** (**a**) For each DNA-bead tether used, at the start of the experiment, the apparent DNA length was recorded at at 60 Hz using *F* = 0.3 pN (raw data, grey) for 5 s intervals at 1 turn increments from 10 to -10 magnet turns (red). The average apparent DNA length was calculated within a window (horizontal bars) to allow for bead settling following each rotation (36). (**b**) A straight-line fit (black) to the average lengths between -6 magnet turns (approximately the R-loop out position) to -4 magnet turns (approximately the R-loop in position) was used as a correction factor to convert DNA extension to a change in turns value. Note that the linear relationship extends below -6 and above -4, as shown.

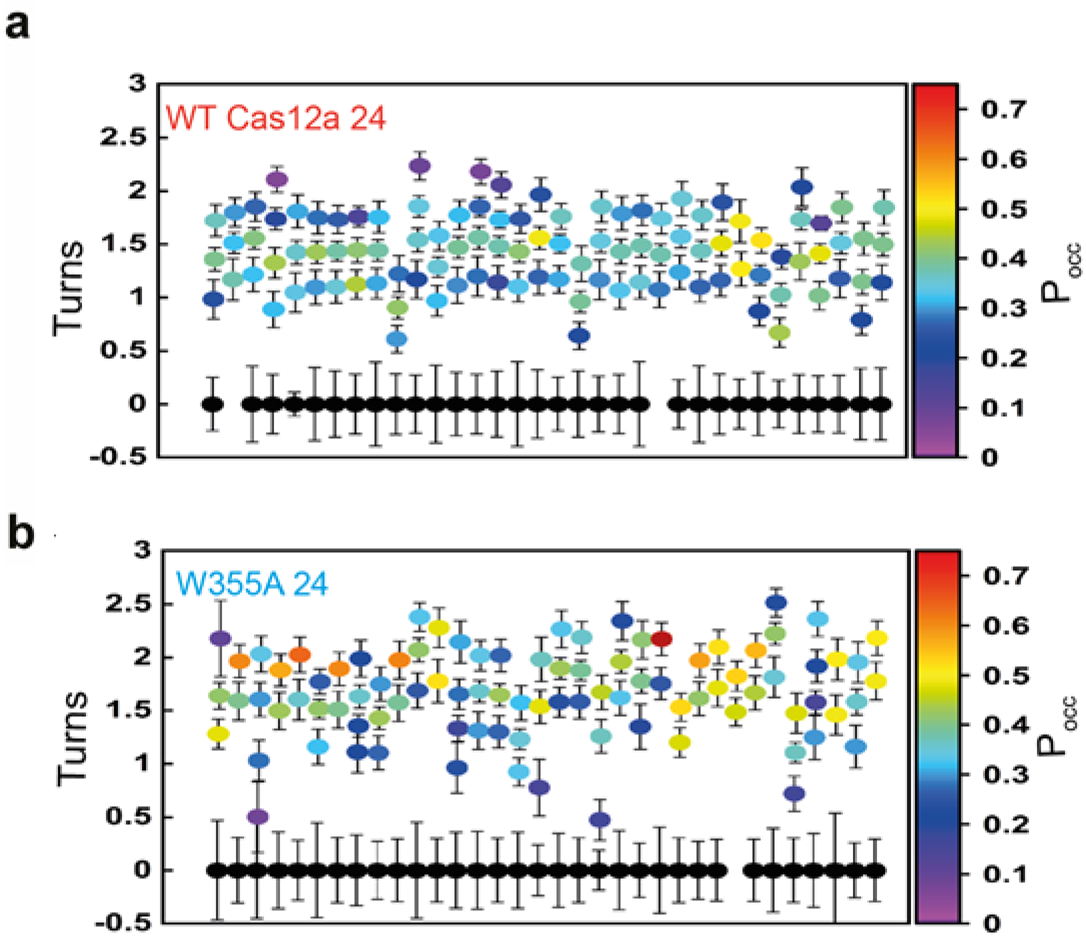


**Supplementary Figure S3. Turns states for individual R-loop events at -7 pN nm.** For each experimental event using 1 nM WT Cas12a (**a**) or W355A Cas12a (**b**) with crRNA 24, individual turn states were identified by HMM (Materials and Methods). The probability of occupancy ($P_{occ}$) of the fitted states was calculated and is given for each state by the heat map. Error bars represent s.d. of the measured turn size. In a few events, the R-loop out state (black) could not be identified as the R-loop formed immediately upon turning to negative torque.

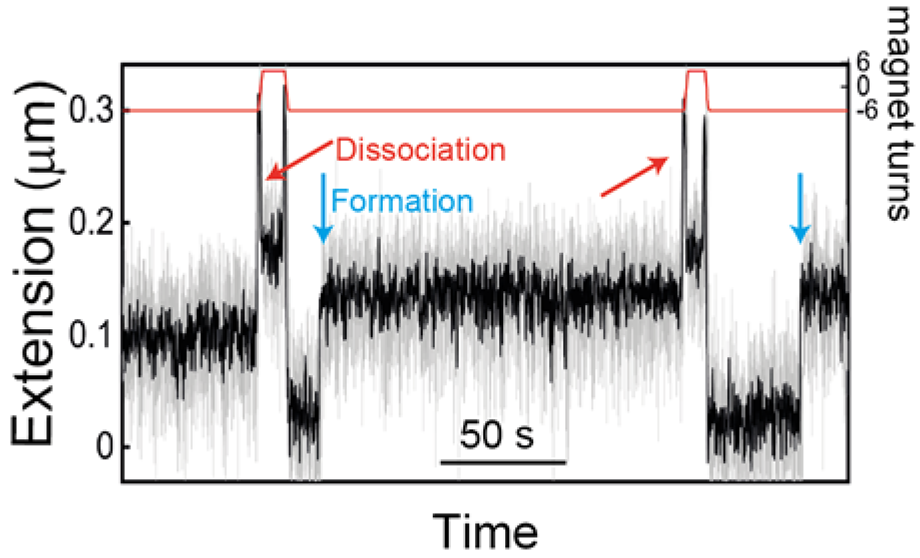


**Supplementary Figure S4. Instability of the W355A Cas12a R-loop state in positive torque.** Bead tracking (light grey, 60 Hz) at 0.3 pN in the presence of 1 nM W355A Cas12a and crRNA 24 is shown filtered to 10 Hz (black line). R-loop formation at negative torque (-6 magnet turns, -7 pN nm) is indicated by the blue arrows. Upon rapidly turning the magnets (10 turns/s) to +4 turns (+7 pN nm), there was immediate R-loop dissociation faster than the time resolution (red arrows).

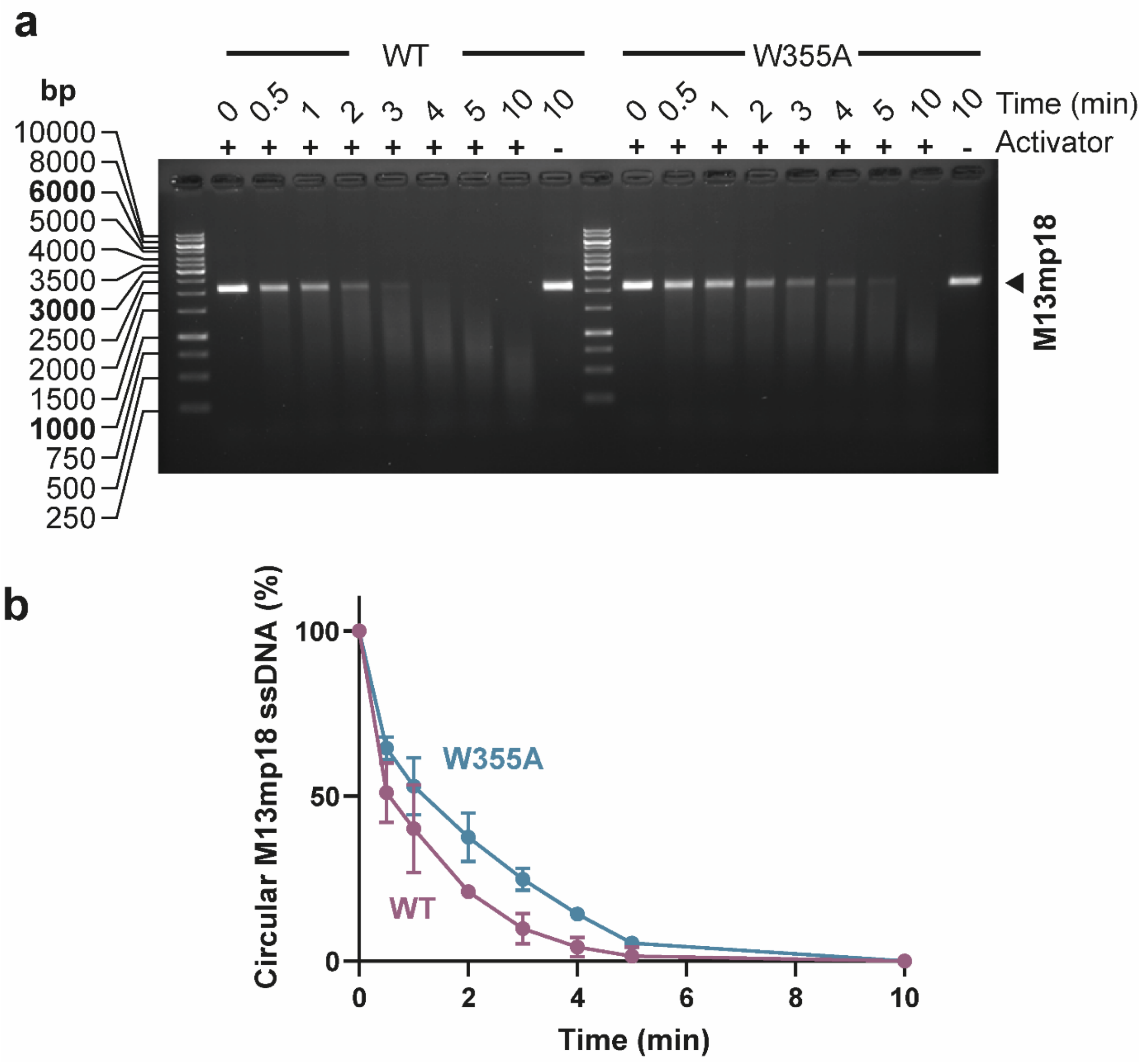


**Supplementary Figure S5. The bystander cleavage activity of WT and W355A Cas12a is similar.** Cas12a-crRNA 24 RNP was preincubated in the presence (+) or absence (-) of a ssDNA TS activator for 30 min. M13mp18 ssDNA was then added, and reaction time points taken at the times shown. Activated bystander activity results in non-specific digestion of the M13 DNA, observed as a smear. The graph quantifies the disappearance of the M13mp18 full length DNA band ($N$ = 3, error bars SD).

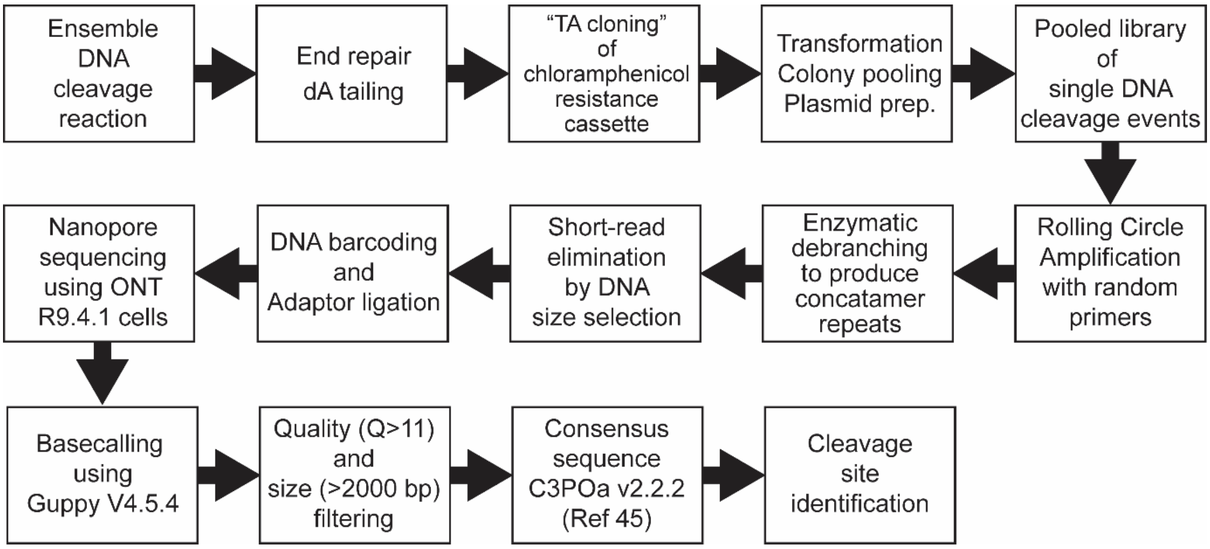


**Supplementary Figure S6. ENDO-pore workflow.** See Materials and Methods for details. Using DNA with ≥5 concatemer repeats produces consensus sequences with >99.5% median accuracy (38).

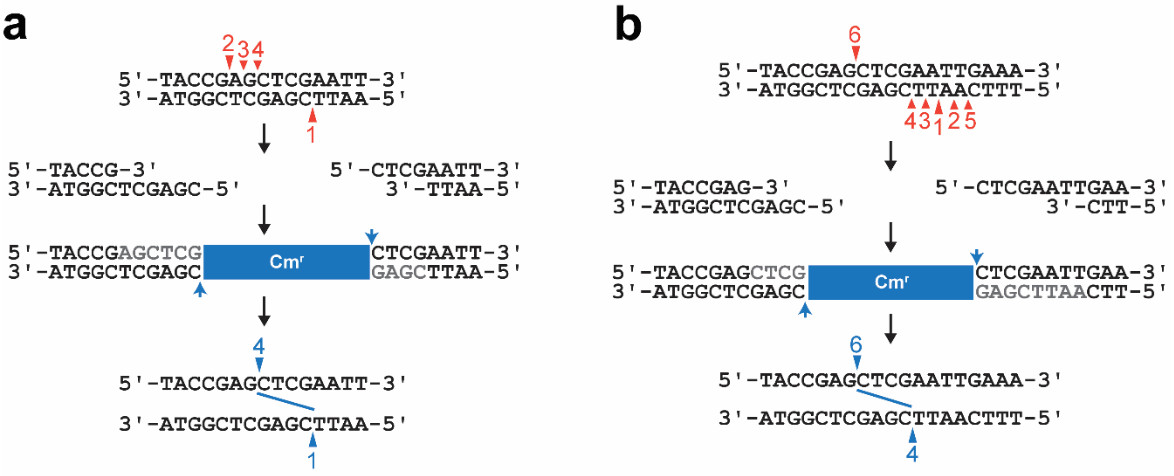


**Supplementary Figure S7. The effect of end repair on the reported sites of DNA cleavage**. When an endonuclease cuts at just one location on the top and on the bottom strand, ENDO-Pore reports the exact cleavage loci (38). Following DNA cleavage and end repair, the ligated chloramphenicol resistance cassette (blue) is used to map the location of the cleavage sites (blue arrows): The 5′ end of the cassette provides the bottom strand cleavage position; the 3′ end of the cassette provides the top strand cleavage position. However, as observed for Cas12a (30), an endonuclease can make secondary cuts that further process the DNA ends. For example: (**a**) The bottom strand is cut at one location (event 1) but there is a subsequent 5′-3′ processing of the top strand (events 2→3→4); (**b**) Following the initial cleavage of the bottom strand (event 1), further random nicking 5′ and 3′ to the break produces a gap (events 2→3→4→5). The top strand is then cut at one location (event 6). In both cases, the cleavage sites reported are those closest to the 3′ ends of the respective strands and are independent of the order of the cleavage events (i.e., panel b).

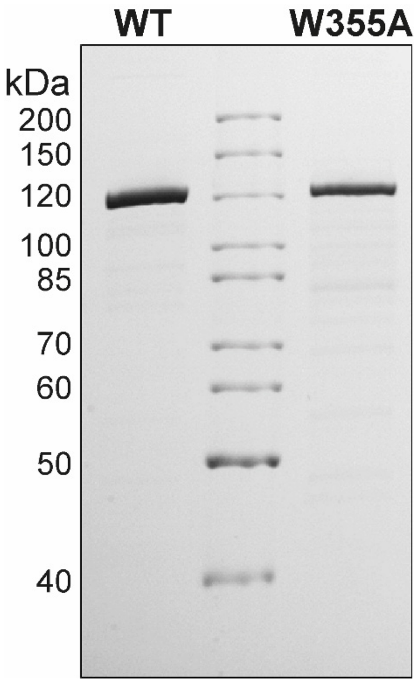


**Supplementary Figure S8. LbCas12a preparations**. 1 µg samples of WT and W355A LbCas12a following purification (Materials and Methods). W355A showed a greater degree of breakdown products.
